## supplemental information for "Multiplexed Assays of Human Disease-relevant Mutations Reveal UTR Dinucleotide Composition as a Major Determinant of RNA Stability"

Supplemental Figure S1

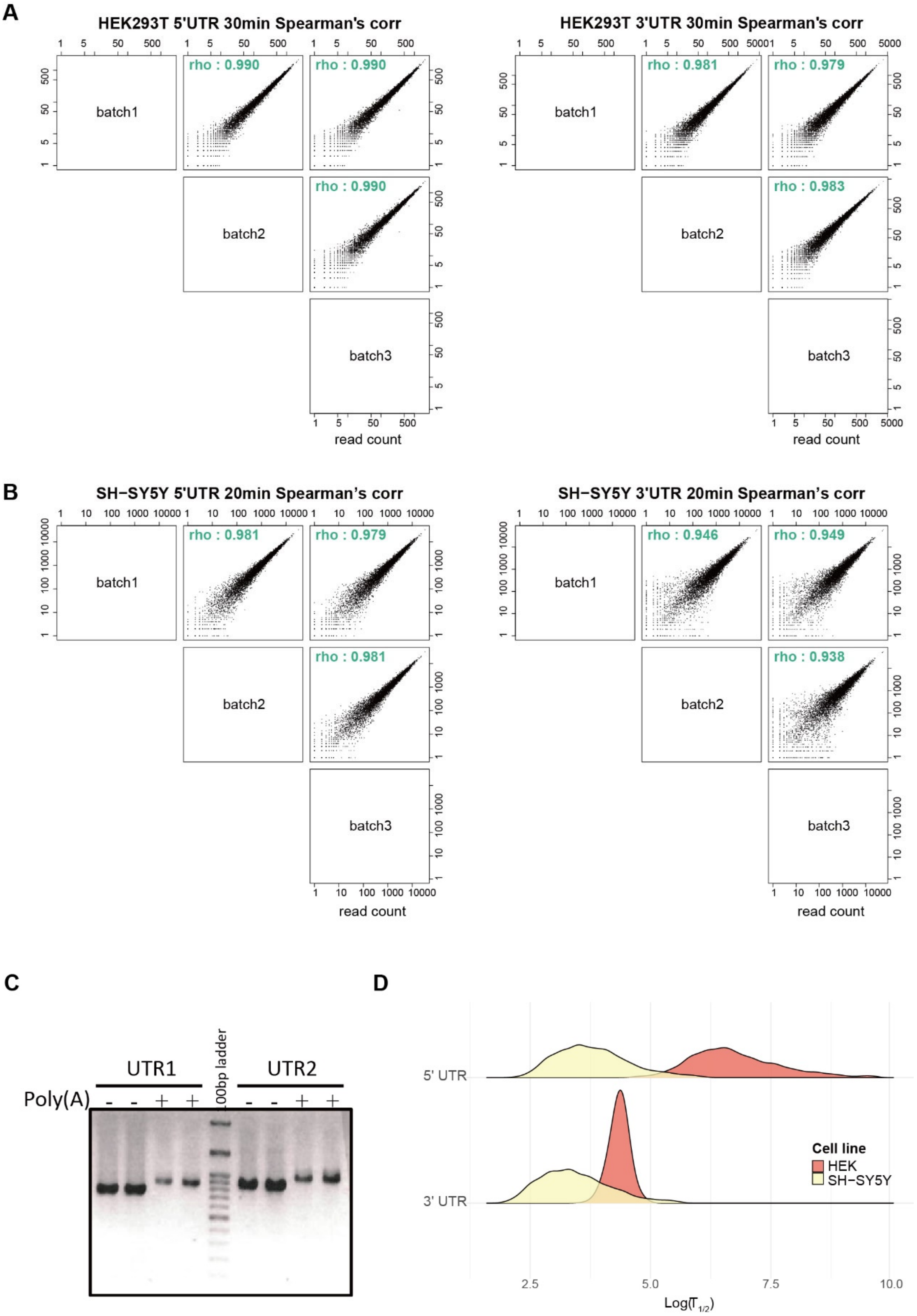


**Supplemental Figure S1. Correlation of experimental results. (A)** Spearman's correlation of the sequencing read counts among three repeated experiments in HEK293T cells. **(B)** Spearman's correlation of the sequencing outcomes among three repeated experiments in SH-SY5Y cells. **(C)** *In vitro* polyadenylation (poly(A)) prior to transfection. **(D)** Comparisons of half-lives in log scale between UTRs and cell lines.

Supplemental Figure S2

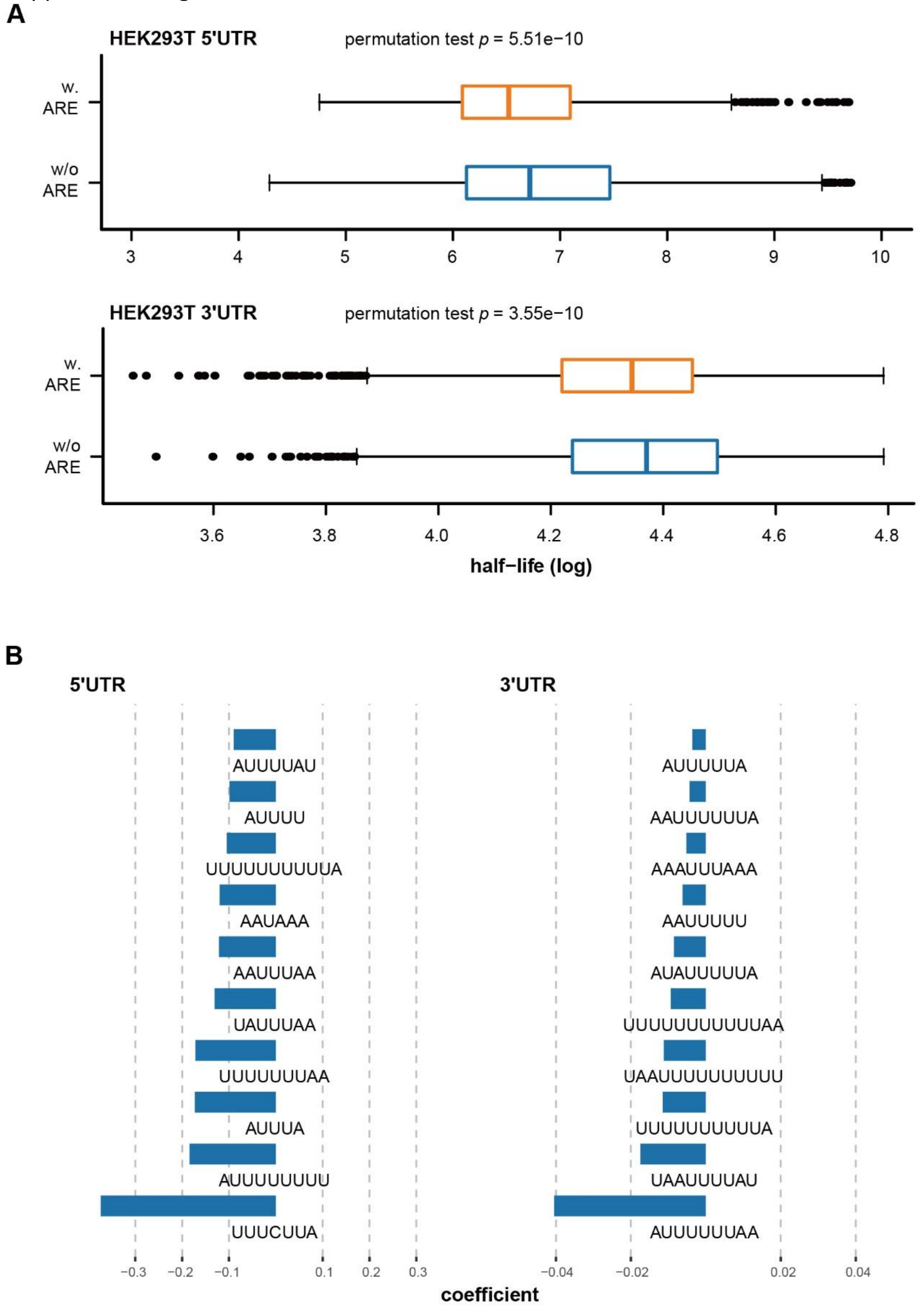


**Supplemental Figure S2. Various destabilizing effects of AREs in HEK293T, related to Figure 2. (A)** AREs of both UTRs destabilize RNA. **(B)** The ten most influential AREs of RNA stability.

Supplemental Figure S3

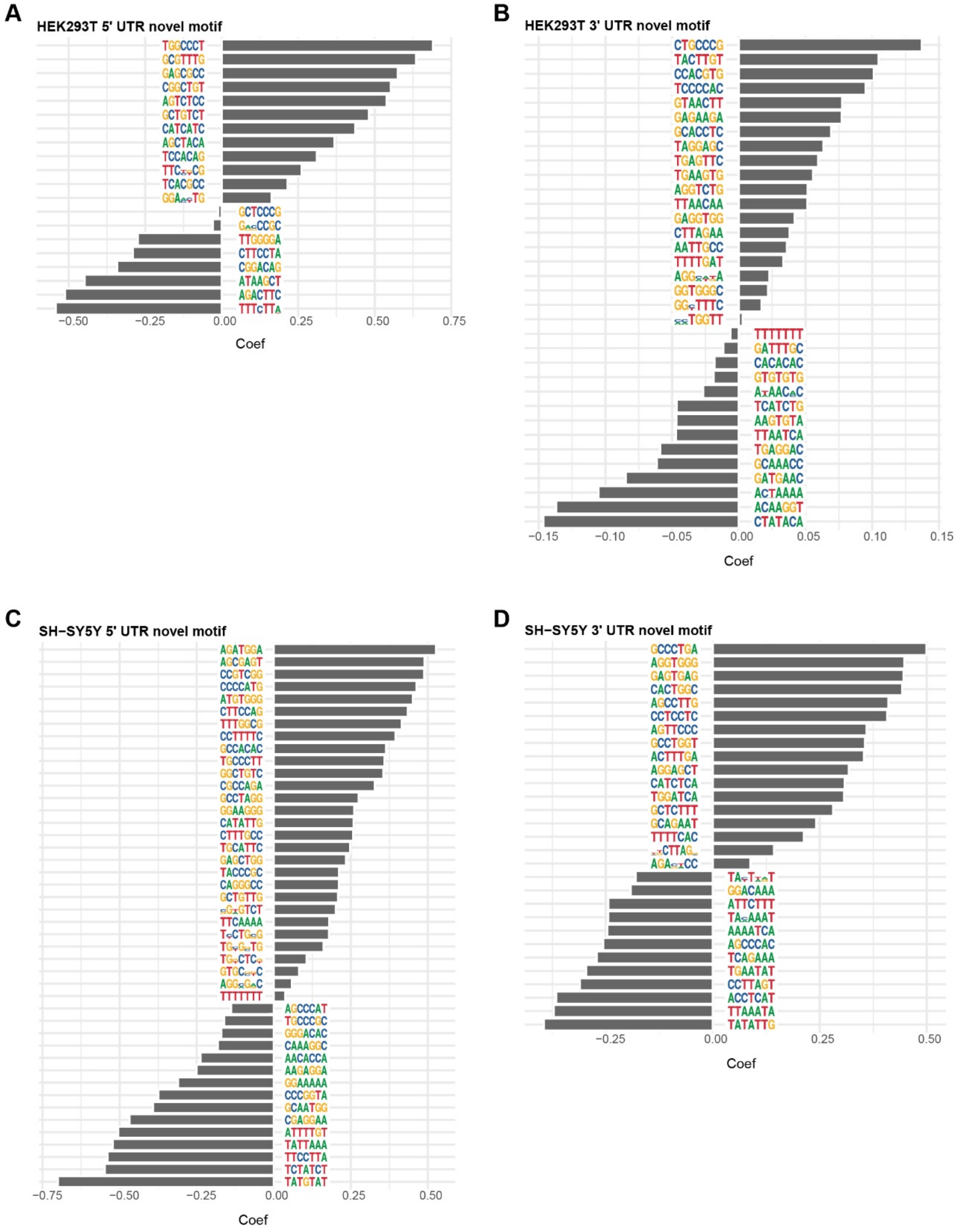


**Supplemental Figure S3. Stabilizing or destabilizing motifs in UTRs.** 7-mer motifs > 20 occurrences in the MPRA library were examined by LASSO regression to test their association with half-life. Motifs that were selected > 1,600 times out of 2,000 bootstraps (see Methods) are presented in this figure: **(A)** HEK293T 5’ UTR motifs. **(B)** HEK293T 3’ UTR motifs. **(C)** SH-SY5Y 5’ UTR motifs. **(D)** SH-SY5Y 3’ UTR motifs.

Supplemental Figure S4

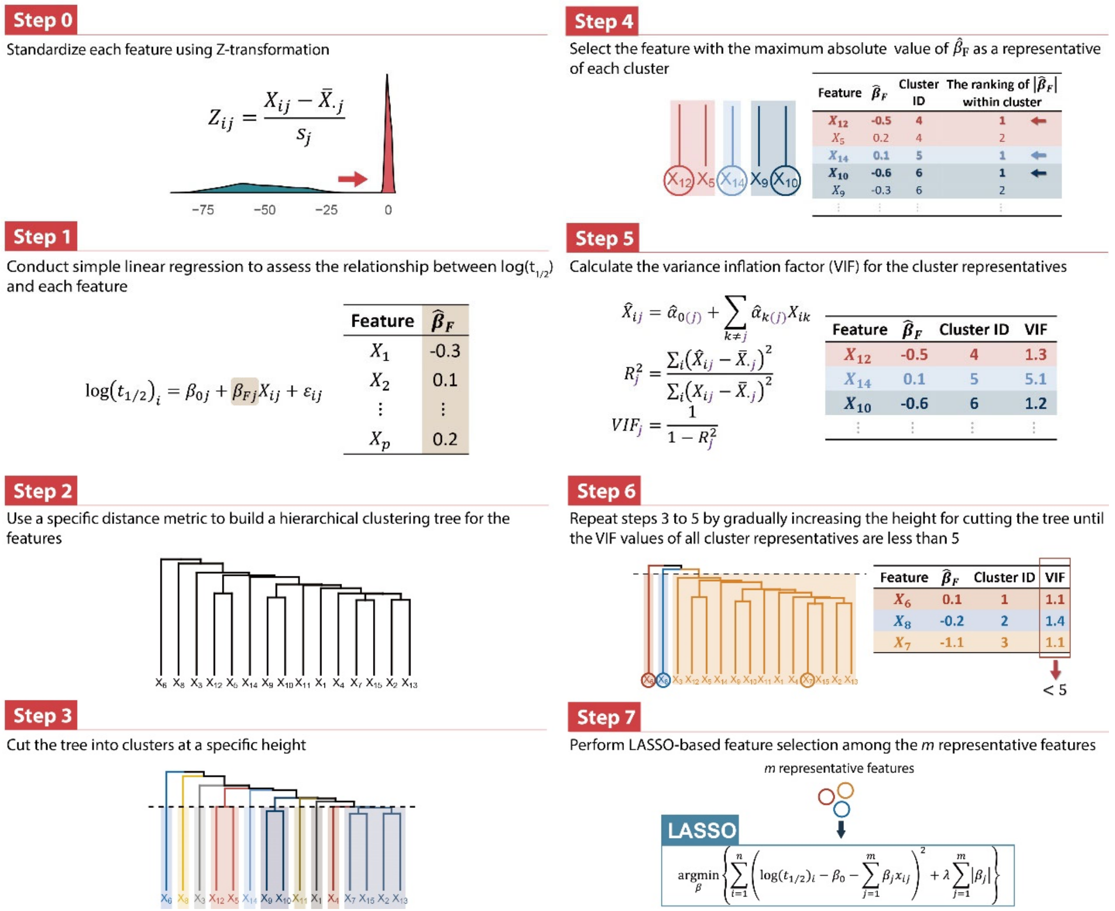


**Supplemental Figure S4. Workflow of variable selection to build models of stability influence.**

Supplemental Figure S5

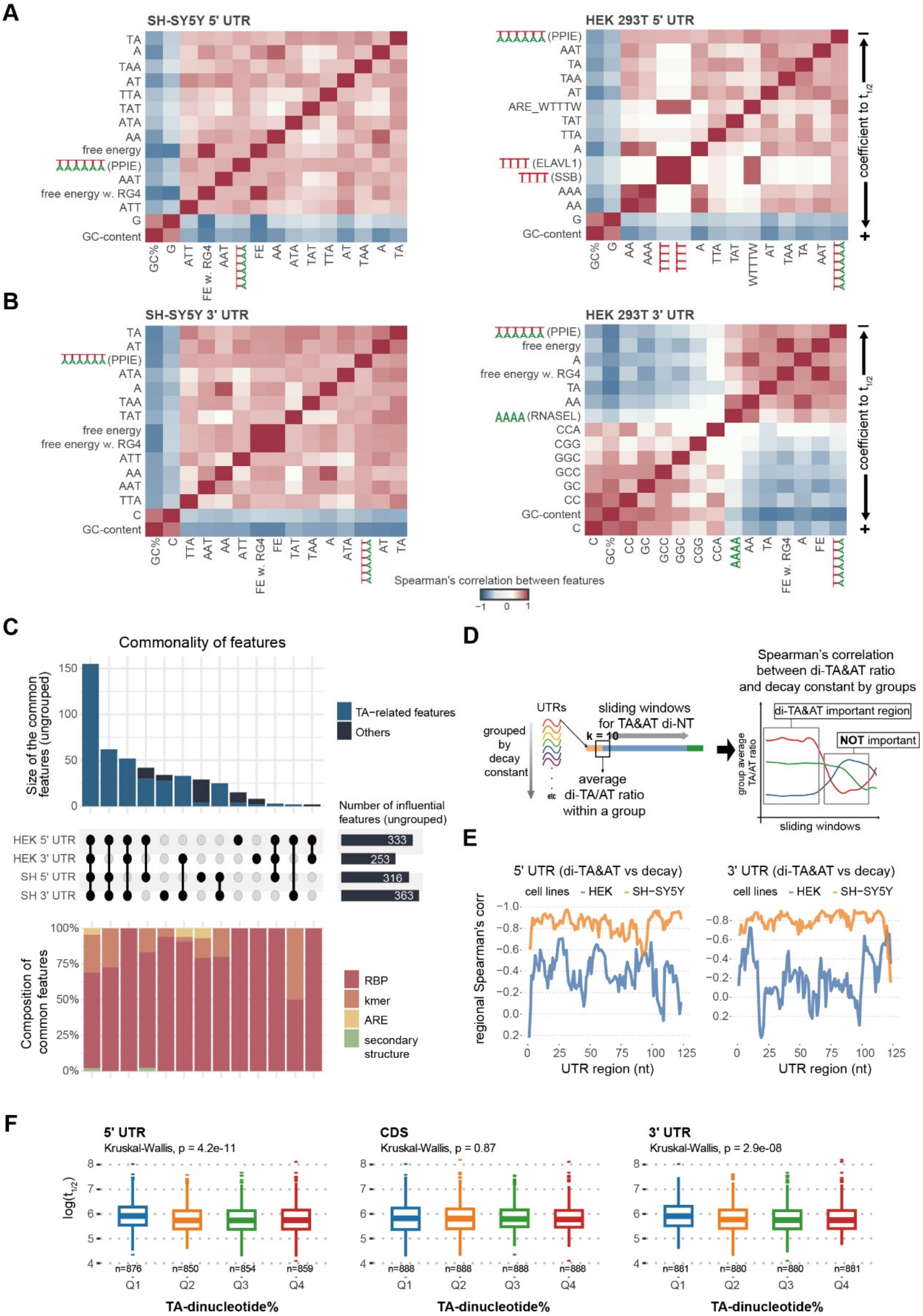


**Supplemental Figure S5. TA cluster is the most potent destabilizing factor in each cell line. (A)** Correlation of top 15 5' UTR influential factors of stability in the TA cluster. **(B)** Correlation of top 15 3' UTR influential factors of stability in the TA cluster, related to Figure 4A-D. Factors are arranged by their coefficient to half-lives. Please note that there are destabilizing factors, such as TA and AT dinucleotides, and stabilizing factors, such as GC-content and G-monomer, in this cluster. TA dinucleotide and WWWWWW (PPIE, W represents A/T) represent the cluster to model UTR stability in SH-SY5Y and HEK293T cells, respectively. **(C)** Commonalities and composition of significant stability-regulating factors among the four experimental groups. **(D)** The workflow of sliding window analysis. A 10-mer window progressing 1 nt at a time calculates the TA/AT dinucleotide ratio within the window. Spearman's correlation between TA/AT dinucleotide ratio and half-lives by groups estimated the importance of regional TA/AT dinucleotide ratio. **(E)** Results of (D) showed a higher correlation between TA/AT dinucleotide and half-lives in SH-SY5Y cells, and a moderate correlation at the end of 3' UTR in HEK293T cells. **(F)** High TA-nucleotide ratios of both UTRs reduce endogenous RNA stability in K562 cells. Q1-Q4 denote quantile groups categorized based on the TA-dinucleotide ratio.

Supplemental Figure S6

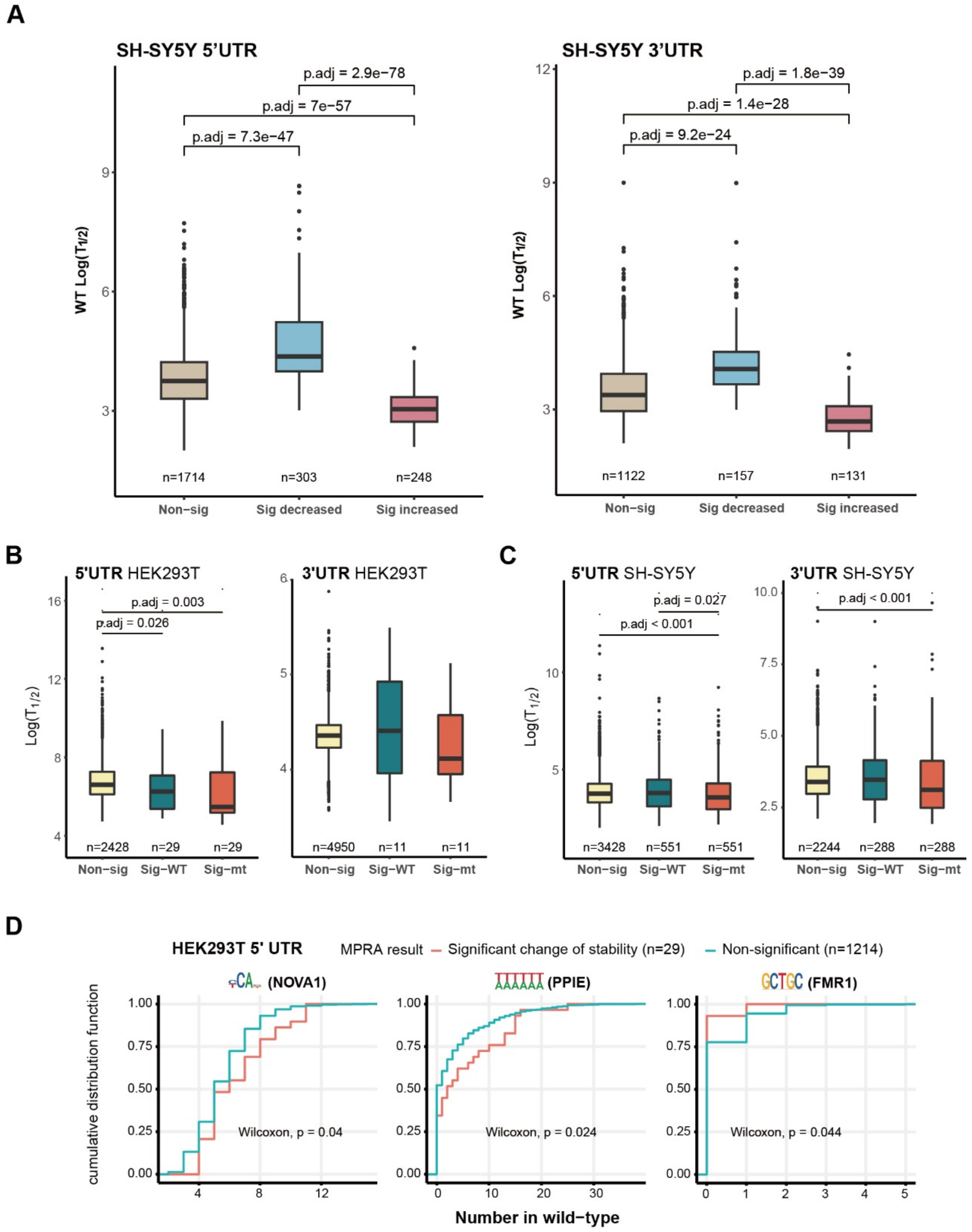


**Supplemental Figure S6. Distinct intrinsic properties associated with stability alteration. (A)** Half-life comparisons among WT UTRs whose mutant counterpart did not alter stability (left), significantly decreased stability (middle) or significantly increased stability (right) in SH-SY5Y cells. Please note that only a few UTR mutations altered stability in HEK293T cells, so this comparison was not done for HEK293T cells due to the limits in sample sizes. **(B-C)** Half-life comparisons among UTRs with no effect from mutations (left), WT UTRs whose mutant counterpart significantly altered stability (middle), mutant UTRs that significantly differed from their WT counterpart in stability (right) in HEK293T (B) or SH-SY5Y cells (C). **(D)** UTRs that are sensitive to mutations (whose mutant altered stability, red lines) are more likely to contain NOVA1, PPIE binding sites, but fewer FMR1 binding sites.

Supplemental Figure S7

**A**

| | SH-SY5Y | | | | HEK293T | | | |
|---|---|---|---|---|---|---|---|---|
| | 5' UTR | | 3' UTR | | 5' UTR | | 3' UTR | |
| | coefficient | *p* value | coefficient | *p* value | coefficient | *p* value | coefficient | *p* value |
| **Univariable** | | | | | | | | |
| $\sqrt{\text{TA}\,\%}$ | -4.55 | $6.3 \times 10^{-47}$ | -4.33 | $6.2 \times 10^{-44}$ | -2.57 | $1.3 \times 10^{-9}$ | -0.29 | $3.1 \times 10^{-6}$ |
| GC content | 1.88 | $7.4 \times 10^{-28}$ | 2.79 | $1.9 \times 10^{-39}$ | 1.27 | $2.1 \times 10^{-8}$ | 0.22 | $3.5 \times 10^{-7}$ |
| **Multivariable** | | | | | | | | |
| $\sqrt{\text{TA}\,\%}$ | -4.68 | $9.0 \times 10^{-21}$ | -2.91 | $1.2 \times 10^{-8}$ | -1.85 | $8.2 \times 10^{-3}$ | -0.08 | $4.7 \times 10^{-1}$ |
| GC content | -0.09 | $7.3 \times 10^{-1}$ | 1.19 | $5.7 \times 10^{-4}$ | 0.48 | $2.0 \times 10^{-1}$ | 0.18 | $3.0 \times 10^{-2}$ |
| **Multivariable with interaction** | | | | | | | | |
| $\sqrt{\text{TA}\,\%}$ | -6.63 | $1.6 \times 10^{-5}$ | -8.90 | $6.4 \times 10^{-7}$ | -6.94 | $4.9 \times 10^{-4}$ | 1.52 | $2.5 \times 10^{-6}$ |
| GC content | -0.82 | $1.8 \times 10^{-1}$ | -1.61 | $6.5 \times 10^{-2}$ | -1.53 | $6.3 \times 10^{-2}$ | 0.98 | $1.2 \times 10^{-8}$ |
| $\sqrt{\text{TA}\,\%}$ × GC content | 3.81 | $1.8 \times 10^{-1}$ | 13.05 | $4.7 \times 10^{-4}$ | 10.67 | $6.2 \times 10^{-3}$ | -3.75 | $1.2 \times 10^{-7}$ |

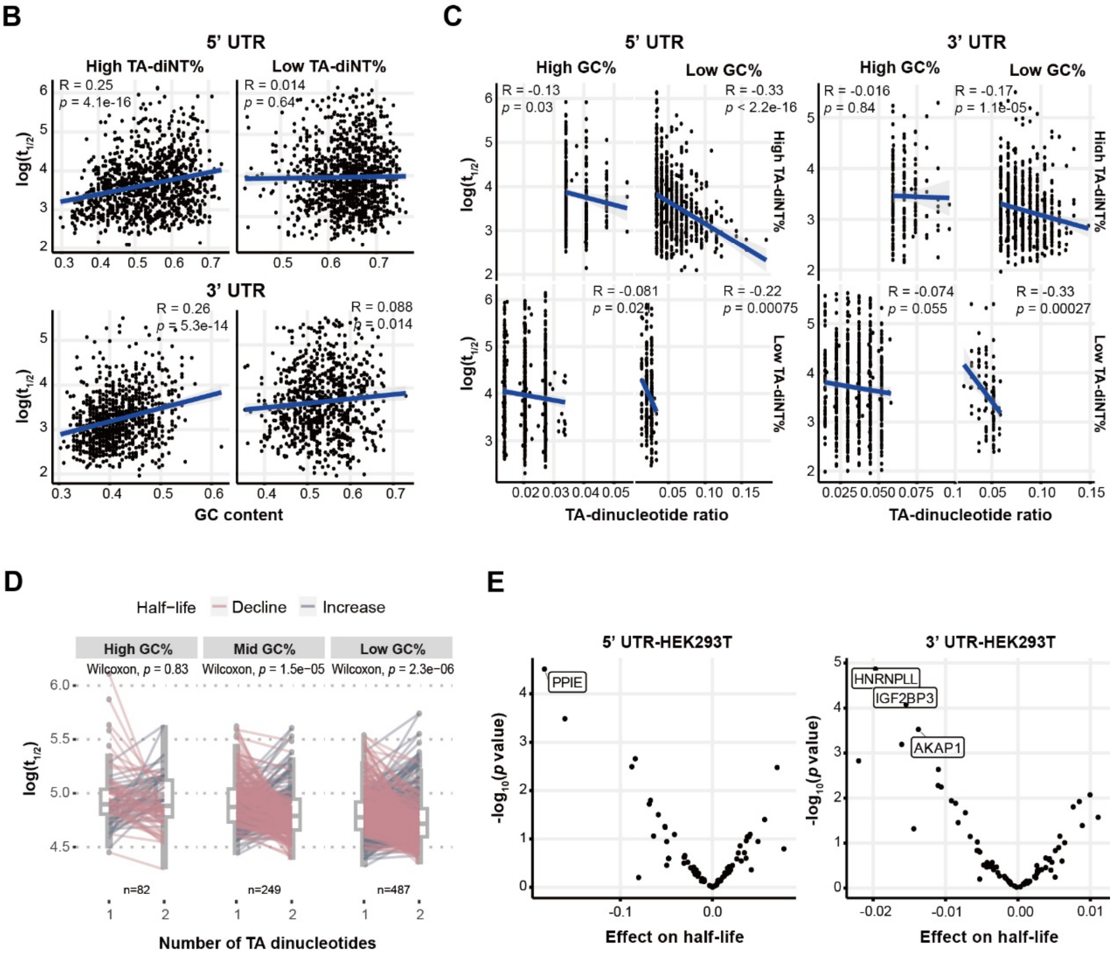


**Supplemental Figure S7. Interplay of GC content and TA dinucleotide on stability regulation, related to Figure 5. (A)** Statistical interaction of GC content and TA dinucleotide on RNA stability. GC content becomes insignificant when considering the effect of TA dinucleotide or TA dinucleotide and their interaction in the multivariate regression. *p* values were determined by linear regression. **(B)** The protective effect of GC content on RNA half-lives depends on the TA dinucleotide ratio. **(C)** Stratifications of both TA dinucleotide ratio and GC content showed that the destabilizing effect of TA dinucleotide is the most prominent under conditions of low TA dinucleotide ratio and low GC content. The same trend was observed for 5' UTR (left) and 3' UTR (right).

**(D)** High GC content mitigates the destabilizing effect caused by the gain of TA dinucleotides in a 5’ UTR random library. **(E)** Association of the binding motifs of TA-dinucleotide-binding protein motifs with RNA half-life in HEK293T cells.
